# Kinetic interplay between droplet maturation and coalescence modulates shape of aged protein condensates

**DOI:** 10.1101/2021.10.07.463530

**Authors:** Adiran Garaizar, Jorge R. Espinosa, Jerelle A. Joseph, Rosana Collepardo-Guevara

**Affiliations:** Maxwell Centre, Cavendish Laboratory, Department of Physics, University of Cambridge, J J Thomson Avenue, Cambridge CB3 0HE, United Kingdom; Yusuf Hamied Department of Chemistry, University of Cambridge, Lensfield Road, Cambridge CB2 1EW, United Kingdom; Department of Genetics, University of Cambridge, Downing Site, Cambridge CB2 3EH, United Kingdom

## Abstract

Biomolecular condensates formed by the process of liquid–liquid phase separation (LLPS) play diverse roles inside cells, from spatiotemporal compartmentalisation to speeding up chemical reactions. Upon maturation, the liquid-like properties of condensates, which underpin their functions, are gradually lost, eventually giving rise to solid-like states with potential pathological implications. Enhancement of inter-protein interactions is one of the main mechanisms suggested to trigger the formation of solid-like condensates. To gain a molecular-level understanding of how the accumulation of stronger interactions among proteins inside condensates affect the kinetic and thermodynamic properties of biomolecular condensates, and their shapes over time, we develop a tailored coarse-grained model of proteins that transition from establishing weak to stronger inter-protein interactions inside condensates. Our simulations reveal that the fast accumulation of strongly binding proteins during the nucleation and growth stages of condensate formation results in aspherical solid-like condensates. In contrast, when strong inter-protein interactions appear only after the equilibrium condensate has been formed, or when they accumulate slowly over time, with respect to the time needed for droplets to fuse and grow, spherical solid-like droplets emerge. By conducting atomistic potential-of-mean-force simulations of NUP-98 peptides—prone to forming inter-protein *β*-sheets—we observe that formation of inter-peptide *β*-sheets increases the strength of the interactions consistently with the loss of liquid-like condensate properties we observe at the coarse-grained level. Overall, our work aids in elucidating fundamental molecular, kinetic, and thermodynamic mechanisms linking the rate of change in protein interaction strength to condensate shape and maturation during ageing.

## Introduction

Living cells contain numerous macromolecular components, which must be organised in space and time to facilitate the concerted regulation of biochemical reactions^1–4^. In eukaryotes, such functional organisation is achieved via the formation of both membrane-bound^5^ and membrane-less compartments^2, 6^. The latter, also termed biomolecular condensates^7^, emerge when, above a critical concentration, key multivalent biomolecules (such as proteins and RNA) spontaneously demix—via the process of liquid–liquid phase separation (LLPS)—into coexisting liquid drops of different compositions^2, 6^. Biomolecular condensates are ubiquitous within both the cytoplasm^8^ and nucleoplasm^9–12^; with the most well-known examples including P granules^13^, nucleoli^14–19^, Cajal bodies^20–22^, or stress granules^23^. Recently, biomolecular condensates have also been identified as functional organisers of the interiors of prokaryotes^24^.

Intracellular LLPS is a delicate phenomenon which is sensitively affected by the environmental conditions (e.g., pH, salt, temperature)^25, 26^, and the presence of different molecular partners^15, 27, 28^. Alteration of such conditions, can lead to misregulation with pathological implications^29–31^. Indeed, the gradual rigidification of biomolecular condensates with time (also known as ‘maturation’ or ‘ageing’) has been associated in the proliferation of multiple neurodegenerative diseases^29–32, 32, 33^—such as amyotrophic lateral sclerosis (ALS)^34^, Parkinson’s^35^, Alzheimer’s^36^, and frontotemporal dementia (FTD)—and in certain types of cancers^37^ and diabetes^38^. Therefore, understanding the molecular mechanisms influencing aberrant LLPS is a key area of research^39^.

Macroscopically, biomolecular condensates present liquid-like properties, such as the ability to coalesce and deform under shear flow^40^, exhibit spherical shapes^2, 29, 41^, show short recovery times from fluorescence recovery after photobleaching (FRAP) or GFP florescence recovery experiments^34, 36, 42^, and exchange material rapidly with their environment^43^. Microscopically, such liquid-like properties originate on the weak multivalent attractive interactions that the biomolecules within the condensate establish. Weak interactions translate into dynamic binding and unbinding, free molecular diffusion within, and facile exchange of species in and out of condensates. Overall, the liquid-like behaviour of molecules enables condensates to fulfill a wide-range of biological functions, from acting as curated reactive volumes that selectively concentrate and exclude specific molecules^4^, buffering of protein concentrations^44^, regulating gene expression^10, 45–47^, sensing changes in the cell environment^48^, to sequestering components harmful in the cell^49^.

Although the liquid-like properties of condensates seem to underpin their functions during health, it is now clear that the material properties of condensates extend far beyond those of low viscous liquids^39, 50^. Indeed, condensates encompass low to high viscosity fluids^51, 52^, hydrogels^53, 54^ and solid-like states^55, 56^. These properties are not surprising if one considers that the physicochemical features of the biomolecules known to form condensates are highly heterogeneous too. These include multidomain, instrinsically disordered, and globular proteins with different chemical makeups^2, 57^, and which can undergo LLPS in pure form via homotypic interactions and/or in partnership with other proteins^15, 58, 59^, RNAs^60–62^, DNA^9, 10, 63^, or chromatin^64–66^ via heterotypic interactions. Furthermore, FRAP, GFP florescence recovery, coalesence, and active and passive microrheology experiments have revealed that over time, even the condensates that start displaying a liquid-like state can ‘mature’ (i.e., change their material properties) and transition to gels or soft glasses^23, 51, 67, 68^. Matured condensates display reduced fusion propensities and longer recovery times after photobleaching^7, 29, 35, 67–72^, which suggest that the diffusion of molecules within is significantly reduced. Several factors have been proposed as key drivers for the liquid-to-solid transition of condensates, including altered salt-concentration or temperature^51, 73^, post-translational modifications^36, 74^, protein mutations^34, 75, 76^, and, most prominently, protein folding and misfolding events^77–82^. All these factors are expected to favour rigidification by increasing the binding affinity among species and slowing down the timescales of inter-protein unbinding events.

In this work, we develop a coarse-grained (CG) simulation approach to investigate the impact of the gradual strengthening of inter-protein interactions—due for instance to the accumulation of inter-protein *β*-sheets^77–79, 82, 83^, post-translational modifications^84^, or changes in the condensate microenvironment^26^— in the the kinetics and stability of protein condensates over time. Our CG simulations reveal that the interplay of the timescales of condensate growth and fusion, and the rate of emergence of stronger inter-protein interactions critically dictates condensate shape: with spherical condensates forming when fusion dominates, and aspherical solid-like states arising when the stronger interactions accumulate faster than the timescales of condensate fusion. Finally, using atomistic simulations, we show that formation of inter-protein *β*-sheets can strengthen interactions sufficiently to trigger the type of dynamical arrest of condensates we observe at the coarse-grained level. Taken together, our simulations provide a time-dependent assessment of the modulation of the dynamic properties of proteins inside condensates, and contrasts kinetics and thermodynamics properties of condensates sustained by strong *vs*. transient inter-protein interactions.

## Results and Discussion

### Strengthening of inter-protein interactions can cause condensate maturation and thermal hysteresis

We begin by investigating how strengthening of inter-protein interactions affects the thermodynamic and rheological properties of condensates. For this purpose, we develop a tailored coarse-grained model that can assess the impact of transient *versus* long-lived protein binding on the kinetic and thermodynamic properties of the condensates they form. Our model approximates an intrinsically disordered protein as a fully flexible Lennard-Jones heteropolymer of 39 beads (i.e., with each bead representing ∼ 8 amino acids) connected by harmonic springs (see Figs. 1A, and section *SIIA* of the Supplementary Information (SI)). Within a single heteropolymer, we combine beads representing both regions prone to establishing stronger interactions (i.e., stickers) with regions that only establish weak interactions (i.e., spacers) (Fig. 1A). Specifically, we distinguish two types of possible interactions among beads: (1) strengthened interactions among amino acids within specific pairs of ‘sticker–sticker’ regions (red–blue beads); (2) weak interactions among all other combinations (blue–blue, red–red, grey–grey, red–grey and blue–grey beads) (Fig. 1A). Pairs of weakly interacting beads (i.e., interaction type 2) interact with a strength equal to *ε*_*D*_, while pairs of beads within the strengthen ‘sticker–sticker’ regions (i.e., interaction type 1) establish interactions 10 times stronger, i.e., equal to *ε*_*S*_ = 10*ε*_*D*_ (Fig. 1A Right). The sequence patterning of the different type of interactions (strong *vs*. weak) of our coarse-grained proteins is shown in Figs. 1A and S2A (Top). Moreover, an alternative patterning for strong *vs*. weak interactions in which beads representing sticker domains are only located along the first half of the coarse-grained sequence (Fig. S2A (Bottom)) is also explored in the *SI* to elucidate possible patterning effects in condensate maturation. The comprehensive description of the coarse-grained potentials and a full list of the model parameters, as well as protein sequences and the employed reduced units are provided in Sections *SIIA* and *SIIB* of the Supplementary Information.

**Figure 1.**
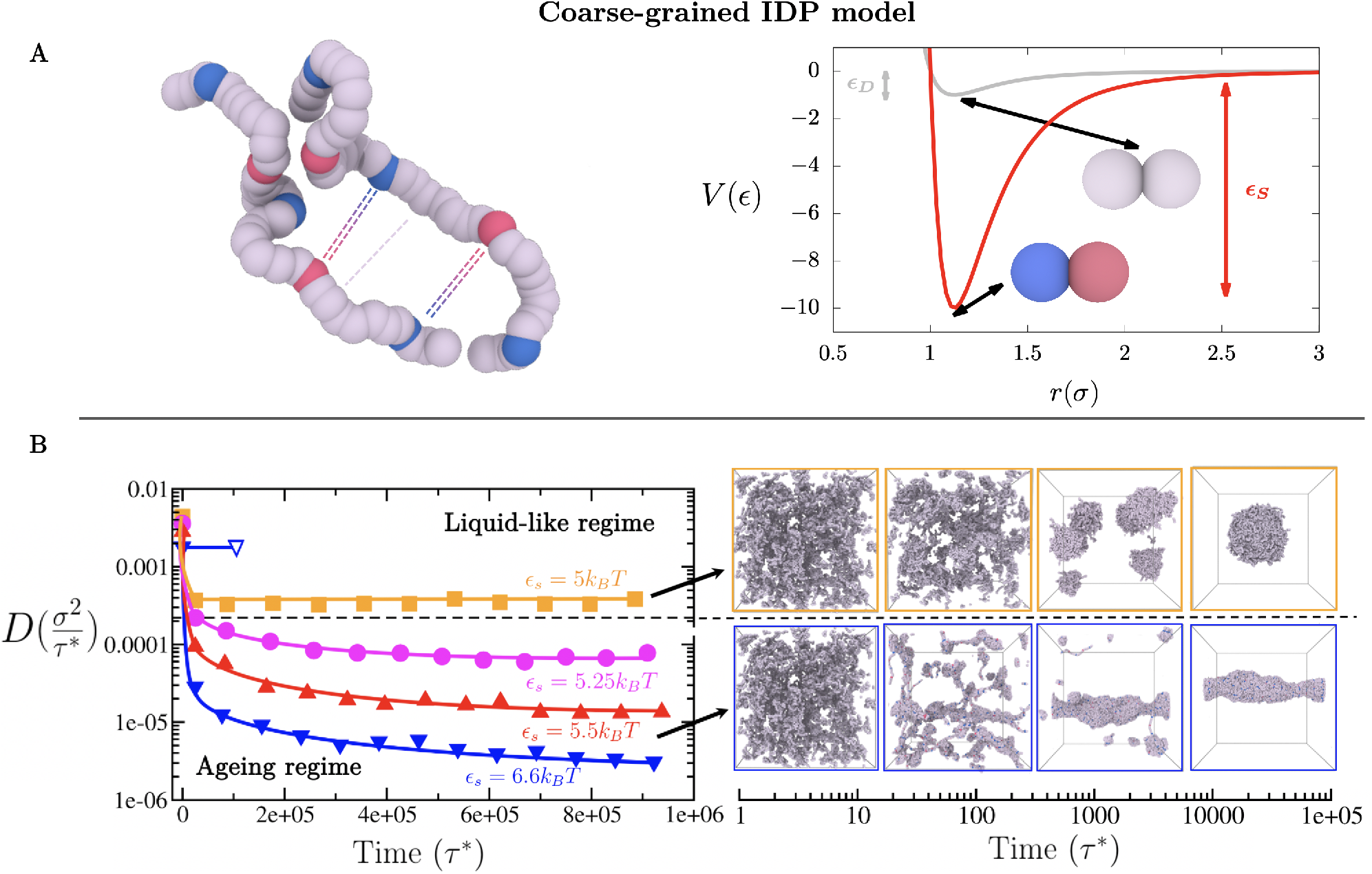
(A) Coarse-grained representation of intrinsically disordered proteins (IDPs) that encodes transient weak interactions between protein domains (grey curve), and strong longer-lived interactions among protein segments (red curve, interactions between blue and red beads). A Lennard-Jones potential of different well-depth (*ε*_*D*_ for interactions between weak interacting domains, and *ε*_*S*_ for interactions between strengthened ‘sticker-sticker’ domains) is used to model the protein binding between different segment types. In our CG model for IDPs, each bead represents a group of ∼ 8 amino acids. Interactions between red and blue beads correspond to *ε*_*S*_, and between all others (grey–grey, blue–grey, red–grey, blue–blue or red–red beads) are weak and transient (*ε*_*D*_). Each protein is composed of 39 beads: 3 blue beads, 2 red beads and 34 grey ones. The excluded volume (*σ*) of each segment type is the same. Results for an alternative CG representation of strong protein binding and a different protein sequence patterning to that depicted in (A) are also available in the Supplementary Information. (B) Time-evolution of the protein diffusion coefficient (*D*) in the condensed phase for different interaction strengths *ε*_*S*_ (in *k*_*B*_T) between strongly-binding protein segments (please note that *ε*_*S*_ = 10*ε*_*D*_). The horizontal black dashed line represents the kinetic threshold of our simulation timescale that distinguishes between ergodic liquid-like behaviour and ageing (transient liquid-to-solid) behaviour. Interaction strengths lower than 5.25*k*_B_T between the strongly-binding segments permit liquid-like behaviour (up to *ε*_*S*_ = 3.5*k*_B_T and *ε*_*D*_ = 0.35*k*_B_T where LLPS is no longer possible), while equal or higher strengths lead to the gradual deceleration of protein mobility over time as shown by *D*. However, in absence of strongly-binding segments (i.e., where all beads bind to one another with uniformly weak binding strength), liquid-like behaviour can be still observed even at *ε*_*D*_ values of 0.66*k*_B_T (empty blue triangle). Black arrows indicate the time dependent behaviour of condensates over time in the liquid-like (Top) and ageing regimes (Bottom). The time evolution snapshots of the condensate corresponds to systems with *ε*_*S*_ = 5*k*_B_T (Top) and *ε*_*S*_ = 6.6*k*_B_T (Bottom). Please note that these snapshots do not correspond to the NVT bulk systems employed to compute the diffusion coefficient in the B left panel.

As a control, we begin by characterising the dynamical properties of peptides inside condensates in the absence of strengthen interactions. In such a homopolymer model, a value of the bead–bead interaction strength, *ε*_*D*_, larger than 0.35 *k*_B_T enables the formation of phase-separated droplets. Therefore, we perform unbiased MD simulations of roughly 750 interacting homopolymers proteins in the *NVT* ensemble, where all beads bind to one another with a uniform binding strength equal to *ε*_*D*_ = 0.66 *k*_B_T; such value of *ε*_*D*_ is high enough to induce condensate formation. From bulk simulations at the equilibrium condensate density, we estimate the mean square displacement (MSD) of the central bead of each protein (in *σ* units, the molecular diameter of every bead in our model), and calculate the value of the diffusion coefficient (*D*) of proteins within the condensates as a function of time (in reduced units *τ*\*) (Empty blue triangle of Fig. 1B; for further details on these calculations see *SIIA* of the Supplementary Information). We observe that the diffusion coefficient of proteins within the droplets (i.e., *D*) quickly converges reaching a value of ∼0.002 *σ*^2^/*τ*\*, characteristic of the free diffusion of polymers within liquids.

We next investigate the change in the mobility of the proteins within condensates when long-lived binding due to strengthening of inter-protein interactions occurs. To do so, we use our heteropolymer model, where 34 beads are treated as spacer regions (i.e., bind to one another weakly with *ε*_*D*_) and 5 beads are treated as sticker regions (i.e., bind to most regions weakly with *ε*_*D*_, but to specific sticker regions strongly with *ε*_*S*_). Note that the value of *ε*_*D*_ controls the strength of interactions among both weakly and strongly binding regions (*ε*_*D*_ = *ε*_*S*_/10). Since the strengthening of inter-protein interactions would depend on the sequence of the amino acids involved^82, 85^ and the physicochemical factors driving such strengthening (e.g. disorder-to-order transitions^77–79, 82, 83^, post-translational modifications^84^, or changes in salt^26^), we explore the dependence of the changes in protein diffusion within condensates on the relative binding interaction strength among beads. Given that values of *ε*_*D*_ larger than 0.35 *k*_B_T enable the formation of phase-separated droplets, we vary *ε*_*D*_ from 0.5 to 0.66 *k*_B_T (Fig. 1B Left). These tests reveal that when proteins bind to one another weakly (*ε*_*D*_=0.5 *k*_B_T, orange curve), the average diffusion coefficient of proteins within the droplets (i.e., *D*) decays moderately (due to the emergence of small clusters of strong inter-protein contacts) and quickly plateaus at a sufficiently high value—signalling ergodic liquid-like behaviour (*D* does not decay over time). In contrast, at stronger protein interaction strengths (*ε*_*D*_ ≥ 0.525 *k*_B_T, magenta, red and blue curves), the diffusion coefficient decays significantly but now fails to reach a plateau within the explored simulation timescale (please note that for measuring *D* over time, we choose the smallest windows of time that allow the central bead of the IDPs to diffuse distances at least 3-5 times their molecular diameter). Such behaviour signals a significant and continuous decay in the protein mobility, consistent with progressive condensate maturation^23, 51, 67, 68^. The reduction in mobility is driven by the gradual formation of strong interactions; this is in contrast to the quick equilibration of the diffusion coefficient in our control simulations, where we treated proteins as weakly-binding homopolymers (even at a value of *ε*_*D*_=0.66 *k*_B_T, blue empty triangle). Decreased mobility of proteins over time, leading to aged condensates (i.e., the ‘ageing regime’), has been inferred experimentally from decelerated diffusion coefficients, higher condensate viscosities^35, 86^ and lower or incomplete recovery from photobleaching^7, 29, 35, 67–72, 87^. Moreover, from Fig. 1B we can observe how the protein diffusion coefficient within the condensates is highly sensitive to small variations in the binding strength between domains. That is, the diffusion coefficient decreases by several orders of magnitude when the binding strength among domains (*ε*_*S*_) is raised from 5 to 6.6 *k*_B_T.

Our simulations reveal that there is a clear inter-protein interaction strength threshold that separates ergodic liquid-like behaviour from non-ergodic ageing behaviour towards glassy droplets (*ε*_*S*_ *>*5 *k*_B_T), which we depict by a horizontal black dashed line in Fig. 1B (Left panel). Above this threshold, condensates readily equilibrate (and form spherical droplets on the simulation timescales), while below it, they gradually become kinetically trapped due to long-lived (i.e., strengthen) interactions that hinder the diffusion of proteins within. Furthermore, the timescale for the onset of strong binding between protein regions during nucleation and growth of the condensates, significantly impacts condensate shape (Fig. 1B Right panel). As expected, condensates that emerge from proteins that bind to one another weakly (i.e., *ε*_*D*_ ≤ 0.5 *k*_B_T) grow into spherical liquid droplets. Spherical shapes are favoured because they minimise the surface to volume ratio and the interfacial free energy cost with the surrounding dilute phase^88^. However, condensates resulting from proteins that bind to one another more strongly (i.e., *ε*_*D*_ ≥ 0.525 *k*_B_T, give rise instead to aspherical kinetically-arrested condensates (Fig. 1B Right Bottom panel). In this case, the emergence of longer-lived interactions prevents individual proteins from relaxing and conveniently rearranging within the condensate to minimise the surface tension, and thus their free energy^89, 90^. We also note that a qualitatively similar behaviour is obtained when the strongly interacting beads are placed along the first half of the sequence (Fig. S3) rather than distributed over its full length (Fig. 1B). Only a moderate increase of the inter-protein interaction strength threshold (*ε*_*D*_ ≥0.6 *k*_B_T) respect to that shown in Fig. 1B (*ε*_*D*_ ≥0.525 *k*_B_T) is required to switch from ergodic liquid-like behaviour to transient ageing behaviour (Fig. S3). Similarly, when the coarse-graining of strengthen interactions is performed through a different potential, as the Wang-Frenkel potential^91^, which significantly reduces the range of strong interactions (Fig. S2B pink curve); we find that just a minor augment of *ε*_*D*_ to values ≥ 0.6 *k*_B_T (and therefore *ε*_*S*_ to ≥ 6 *k*_B_T) is needed to bring condensates from a liquid-like state into the ageing regime originating the emergence of amorphous kinetically arrested droplets (Fig. S5). Hence, the observed behaviour in Fig. 1B seems to be qualitatively independent of moderate variations in the patterning of strong *versus* weak interactions or to distinct coarse-grained approaches to mimic strong protein binding.

We also study the thermal hysteresis of matured condensates. To that end, we set a protein interaction strength that enables phase separation at T=300 K (*ε*_*D*_=0.5 *k*_B_*T*; Top panel in Fig. 2A). From the initial homogeneous system, proteins nucleate several small spherical droplets that grow and coalesce, eventually yielding a single spherical larger condensate (i.e., the global free energy minimum in the liquid-like regime). By starting from the same homogeneous system, we then increase the interaction strength among proteins to *ε*_*D*_=0.66 *k*_B_*T* (ageing regime) in order to promote the formation of stronger longer-lived interactions. As expected from Fig. 1B, we observe the emergence of an elongated kinetically-arrested condensate (Fig. 2A (Left bottom panel)). Interestingly, when reducing the protein interaction strength back to the previous value (*ε*_*D*_=0.5 *k*_B_*T*), thereby mimicking the heating of an aged condensate, the aggregate remains in the dynamically arrested configuration, instead of relaxing into a spherical liquid-like condensate as in Fig. 2A (Top panel). Accordingly, the system displays hysteresis behaviour. We note that hysteresis will widely depend on the associated binding energy of the strongly binding protein segments, and on the thermodynamic pathway. Therefore, subtle variations on these magnitudes (i.e., in our model *ε*_*S*_ and *ε*_*D*_) can significantly impact the degree of hysteresis^92^. For the sequence patterning with regions that can establish strong bonds only distributed over the first half of the chain (Fig. S2A (Bottom)), thermal hysteresis can be observed within a similar range of temperatures (Fig. S4). However, when stronger interactions are modelled through the Wang-Frenkel potential^91^ but maintaining the original sequence patterning (Figure 1A), hysteresis is only observed until *ε*_*D*_ ∼ 0.58 *k*_B_T (Fig. S6). These results highlight how small differences in sequence patterning (Fig. S4) or intermolecular interaction range (Fig. S6) can lead to moderate variations in the thermodynamic pathway and conditions arresting condensate dynamics and leading to thermal hysteresis. Nevertheless, we have also verified for all our simulations that when heating up the matured condensate even further (by changing *ε*_*D*_ from 0.66 *k*_B_*T* to 0.35 *k*_B_*T*, a value too weak to sustain LLPS), full dissolution occurs. Such behaviour correctly recapitulates the moderate thermal hysteresis of hydrogels sustained by reversible fibrils^77–79^ or the salt resistance hysteresis observed in different RNA-binding proteins such as FUS^51^ or hnRNPA1^67, 82^.

**Figure 2.**
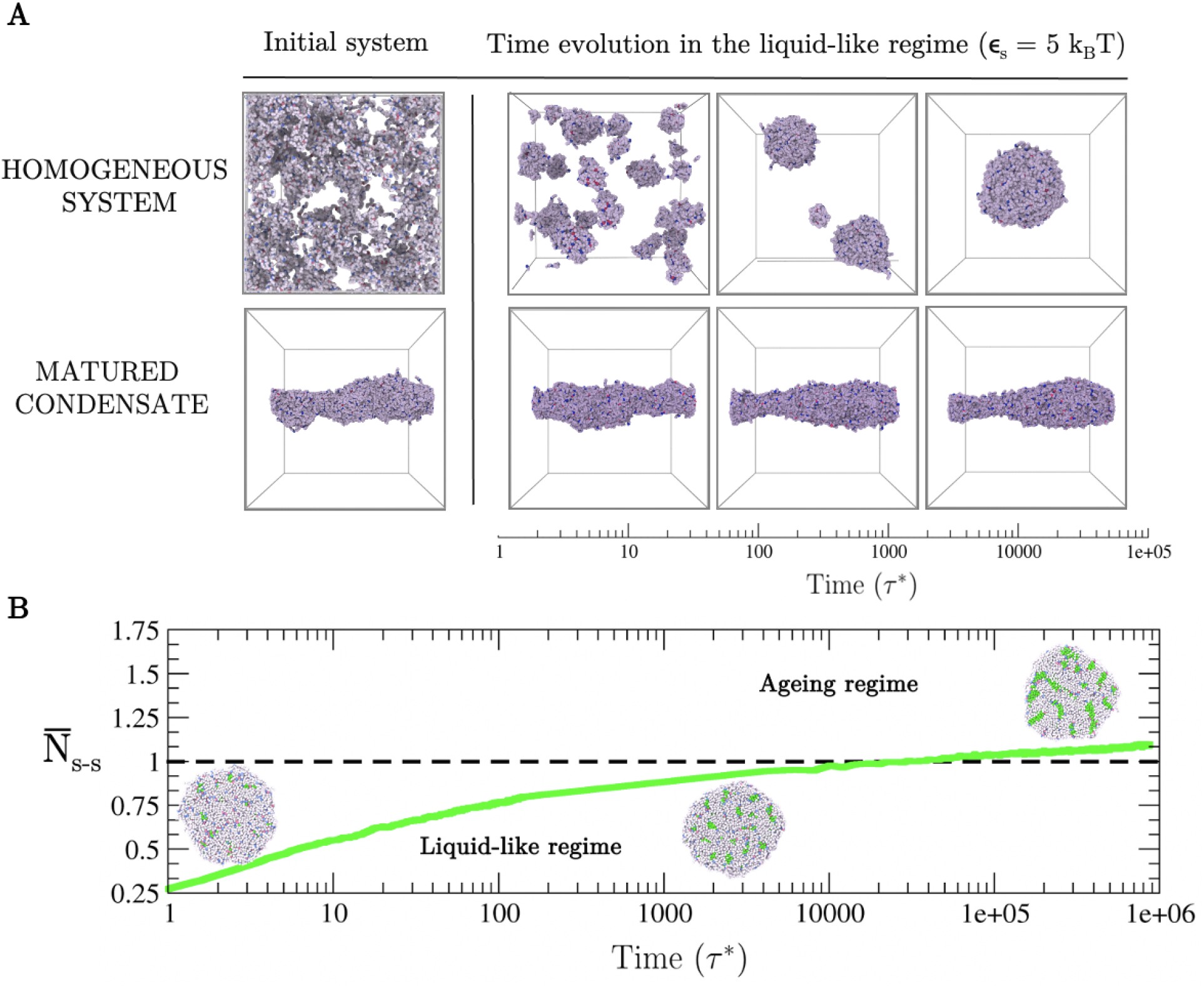
(A) Thermal hysteresis of the condensates probed via coarse-grained protein simulations. (Top panel) Time-evolution starting from an homogeneous system where inter-protein interactions are moderate (i.e., *ε*_*S*_ = 5*k*_B_*T*; *ε*_*D*_ = 0.5*k*_B_*T*). (Bottom panel) Time evolution at the same conditions above, although starting from a matured condensate that was formed under ageing regime conditions (i.e., strong inter-protein interactions of *ε*_*S*_ = 6.6*k*_B_*T*). Note that in our model, temperature *T* is proportional to 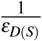. (B) Number of strong interactions as a function of time within a preformed spherical condensate 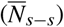 normalised by the typical strong contact threshold (horizontal dashed line) that induces ageing behaviour of protein condensates at those conditions (i.e., number of strong interactions per condensate volume found at the cross-over of the blue curve with the kinetic threshold shown in Fig. 1B). Snapshots of the condensate shape as a function of time are shown. Protein segments that do not participate in strengthen contacts are depicted in grey, while those involved in clusters of stronger interactions are coloured in green. The protein interaction strength of this simulation was set to *ε*_*S*_ = 6.6*k*_B_T, the same set value for the condensates shown in the bottom left panel of Fig. 1B

Strikingly, we also find that when stronger inter-protein interactions emerge inside already formed spherical droplets, instead of during the condensate nucleation and growth stages, the droplets only experience a modest shape deformation over time (Fig. 2B). To interrogate this behaviour, we estimate the average number of strong inter-protein contacts required to drive the condensate out of the liquid-like regime to a kinetically arrested state (i.e., strong contacts per condensate volume at the cross-over between liquid-like and ageing regimes in Figure 1B). We then assess the number of strong inter-protein interactions as a function of time (green spheres in Fig. 2B, further details on these calculations are provided in Section *SIIC*), and observe that even after crossing the threshold number of strong inter-protein interactions needed to trigger ageing behaviour (horizontal dashed line), condensates still remain roughly spherical. This behaviour is likely due to the extremely low diffusion of proteins that are engaged in the long-lived interactions. Although such proteins can rearrange locally to maximise their enthalpic gain via strong inter-protein bonds, they cannot diffuse sufficiently to yield alternative condensate arrangements of potentially lower free energy. In that respect, our results are only consistent with the widely recognised asphericity as a consequence of condensate maturation^35, 51^, as long as the origin of such asphericity is mainly due to non-ergodic droplet coalescence, as discussed in the following section. Fusion of small protein clusters to matured condensates is expected to significantly contribute to the formation of aspherical condensates as shown in Fig. 1B (Bottom Right panel). Moreover, impaired exchange of molecules between condensates and their surroundings, as observed in different multivalent proteins^62, 93^, can lead to the emergence of irregular morphologies. In that respect, mean-field model simulations have been shown to be extremely useful in providing thermodynamic guidance on hardening phenomena^94, 95^.

### Kinetic interplay between droplet maturation and coalescence

Finally, we investigate the kinetic competition between condensate maturation and growth due to droplet coalescence. Accordingly, we evaluate the time required for complete coalescence (*τ*_*c*_), which we term coalescence time, of two spherical droplets (in tangent contact) into a single spherical condensate. We calculate the coalescence time for pairs of droplets of different sizes (from 50 to 500 proteins each) and for distinct values of strong inter-protein binding strength (Fig. 3). We observe that weak inter-protein interaction strengths allow complete fusion of the two initial droplets into a single larger spherical droplet (filled squares; liquid-like regime). Moreover, small droplets fuse considerably quicker than large condensates (i.e., 50-protein droplets fuse up to four times faster than 500-protein condensates at *ε*_*S*_ = 5*k*_B_*T*). However, as the inter-protein interaction strength increases—for instance due to the emergence of solid-like nuclei due to the enhanced binding of proteins—and/or the coalescence time slows down, the initial condensate pairs are no longer able to rearrange into a single spherical droplet on the accessible simulation timescales due to condensate maturation (empty squares; ageing regime).

**Figure 3.**
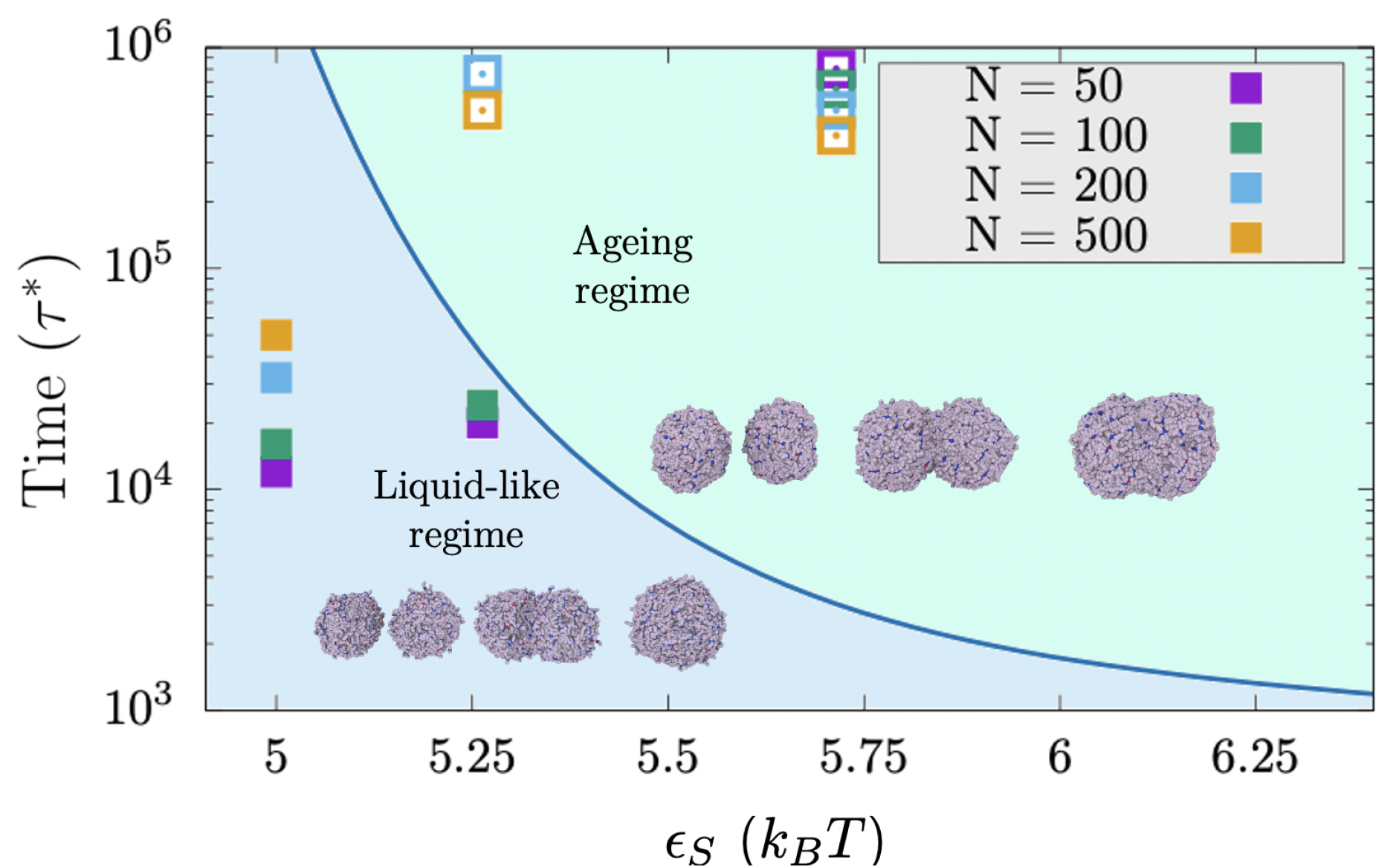
Competition between droplet coalescence time and maturation rate as a function of strong inter-protein interactions (*ε*_*S*_) for different droplet sizes. Blue curve depicts the lapse of time before proteins enter into the ageing (kinetically-trapped) regime due to the emergence of long-lived contacts. The blue curve is a kinetic line that is defined as the intersection of the different diffusion curves and the horizontal kinetic threshold shown in Fig. 1Bs (Left panel). Filled squares represent the time required for two spherical tangent droplets of a given size to fuse into a single spherical condensate, while empty squares depict the maximum simulated time for tangent droplets that did not achieve complete coalescence into a spherical shape. Snapshots of the typical time-evolution of coalescing droplets in both regimes at *ε*_*S*_ = 5.25*k*_B_T are included for droplet sizes of 100 (liquid-like regime) and 200 proteins (ageing regime).

To highlight how the interplay between coalescence time and protein interaction strength commits condensates to either the liquid-like or ageing regime, we define a kinetic threshold (blue curve in Fig. 3) from the intersection between the diffusion coefficient curves and the horizontal black dashed line shown in Fig. 1B (Left panel). This curve should allow us to (a priori) separate the liquid-like region from the ageing regime. At weak inter-protein interactions, coalescence needs to be extremely slow for condensates to enter into the ageing regime before complete fusion takes place, while at strong inter-protein interactions even fast coalescence times may result in aged aspherical droplets. Importantly, we find that the inferred threshold from bulk protein diffusivities in Fig. 1B effectively discriminates the behaviour of kinetically trapped condensates (empty squares) from those that can equilibrate into spherical droplets (filled squares) in our coalescence simulations (Fig. 3).

These results further demonstrate how droplet shape can be critically modulated by the competition between two distinct timescales: coalescence time and maturation rate. This behaviour is particularly well exemplified by the simulations at inter-protein interaction strengths only just sufficiently high to give rise to arrested glass-like behaviour (i.e., *ε*_*S*_ ≥ 5.25 *k*_B_ T). In this case, for the smallest droplet sizes (50 and 100 proteins per droplet, purple and green squares respectively in Fig. 3), the time required for droplet fusion and minimisation of the system’s free energy (i.e., forming a spherical condensate) is shorter than the maturation time (i.e., the formation of long-lived interactions that cause the system to become kinetically trapped). However, for larger droplet sizes (i.e., those containing 200 and 500 proteins), the condensate (both tangent droplets) becomes kinetically arrested before achieving a spherical arrangement. On the other hand, moderate inter-protein interactions permit the complete coalescence of all tested condensate sizes into spherical droplets (*ε*_*S*_ = 5 *k*_B_ T), while stronger interactions (i.e., *ε*_*S*_ = 5.75 *k*_B_ T) do not yield complete fusion of even the smallest tested droplets (Fig. 3). In that respect, our simulations highlight how small variations in the binding energy between protein domains can crucially modulate the liquid-like behaviour, and ultimately the shape of biomolecular condensates.

Based on Figs. 2 and 3, we argue that condensate asphericity seems to be fundamentally determined by fusion events of kinetically arrested droplets rather than from maturation of preformed spherical condensates. Moreover, we note that different patterning^96^ of strong-binding domains along the protein sequence (Figures S3 and S4), does not show a qualitatively distinct behaviour to that of Fig. 2 in terms of shape evolution along condensate maturation^39^.

### Comparing the interaction strength of inter-protein interactions among disordered *vs* ordered peptides

In the section, we quantify the change in the strength of inter-protein interactions due to the formation of inter-protein *β*-sheets to determine if such change may be consistent with the dynamical arrest we describe in our coarse-grained simulations. We are particularly interested in the formation of inter-protein *β*-sheets because they can emerge spontaneously and intrinsically, i.e., without requiring changes in the chemistry of the system or the environmental conditions. Interestingly, the intrinsically disordered regions of various phase-separating naturally occurring proteins—including fused in sarcoma (FUS)^78^, TAR DNA-binding Protein of 43 kDa (TDP-43)^79^, heterogeneous nuclear ribonucleoprotein A1 (hnRNPA1)^77, 82, 83^, and nucleoprotein of 98 kDa (NUP-98)^77, 97^—which form hydrogels over time^76, 98, 99^, contain short regions termed Low-complexity Aromatic-Rich Kinked Segments (LARKS) that are prone to form such inter-protein *β*-sheets^85^. When multiple LARKS meet at the high concentrations found inside condensates, they can assemble into ordered arrays of inter-protein *β*-sheet structures that stick to one another strongly via *π*–*π* bonds and hydrogen bonding between backbone atoms that may lead to gradual solidification of, otherwise, liquid-like condensates^75, 77, 80, 89, 100, 101^. Importantly, hundreds of protein sequences capable of such disorder-to-order conformational transitions, and concomitant enhancement of intermolecular binding strengths, have been identified in the human genome^77^.

As a case study, we focus on the NUP-98 protein—an aggregation-prone protein that phase separates *in vitro* under selective conditions and can form hydrogels under others^102, 103^. We start by estimating the binding strength among four interacting NUP-98 LARKS-containing peptides by means of Umbrella Sampling Molecular Dynamics simulations^104^ in explicit solvent and ions under two distinct scenarios: (1) when all the peptides are fully disordered, and (2) when peptides form the inter-peptide cross-*β*-sheet motif resolved crystallographically (PDB code: 6BZM)^77^. From these simulations (using the a99SB-*disp* force field^105^), we compute the potential of mean force (PMF) as a function of the centre-of-mass (COM) distance between one single peptide—which we gradually force to dissociate from the other segments—and the other three segments (simulation details are described in section *SI* of the Supplementary Information). For the scenario when LARKS are treated as fully disordered, we allow peptides to freely sample their conformational space (only fixing the position (in the appropriate direction) of the closest atom to the peptide COM of the structured four-peptide array; see SI for further details). In the second scenario, where we quantify the interactions among ordered LARKS, we constrain the peptides to retain their crystal *β*-sheet structure^77^.

Our simulations show that the interaction strength between disordered unconstrained peptides is sufficiently weak (i.e., < 0.5*k*_B_T per residue) that, at room temperature, thermal fluctuations would frequently break and re-form such inter-protein interactions, consistent with the formation of liquid-like condensates (Fig. 4, grey curve). More interestingly, when the peptides assemble into constrained inter-peptide cross-*β*-sheet structures, the strength of their interactions increases by almost an order of magnitude (i.e., to approximately 4*k*_B_T per residue, red curve). To verify that our conclusions on the relative difference between disordered and structured binding are not model dependent, we also compute the PMF dissociation curve using the CHARMM36m force field^107^. As shown in Fig. S1 of the SI, a ten fold difference between both peptide dissociation curves is also obtained in agreement with our calculations using the a99SB-*disp* force field^105^ (Fig. 4). We note, however, that the exact magnitude of this increase may be slightly overestimated by the constraints we have used to enforce the stability of *β*-sheet structures. Nevertheless, these results, together with our coarse-grained simulations, suggest that an enhancement of inter-protein interactions may occur due to the formation of inter-peptide LARKS *β*-sheets, sufficient to sustain the formation of gels or aged solid-like aggregates. Importantly, the strength of structured LARKS–LARKS interactions is sufficiently weak that they can still be considered thermolabile. Our results are also consistent with experiments reporting that LARKS-containing proteins form reversible hydrogels that can be easily dissolved with heat^77–79^. A significant increase in the interaction strength after a disorder-to-order transition has been reported previously for the *Aβ* 1–42 system^80^. However, in the case of *Aβ* 1–42, the observed increase was much larger, consistent with amyloid fibers being thermostable^108, 109^.

**Figure 4.**
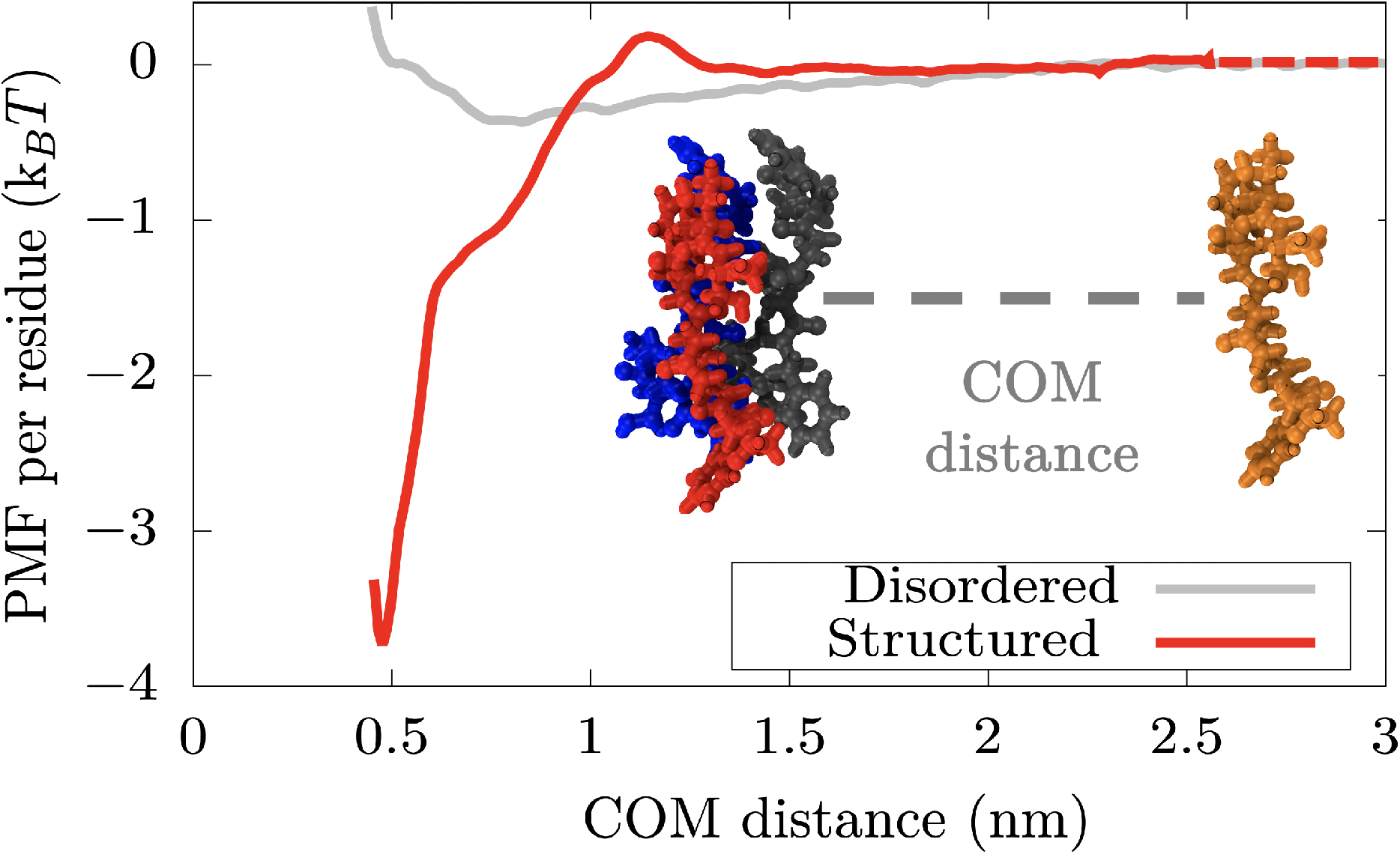
(A) Atomistic Potential-of-Mean-Force (PMF) dissociation curve of an 8-amino acid segment (PDB code: 6BZM) of NUP-98 protein from a *β*-sheet structure of 4 peptides (of the same sequence) as a function of the center of mass distance (COM) using the a99SB-*disp* force field^105^. Red curve depicts the interaction strength among peptides with a well-defined folded structure, kinked *β*-sheet structure, while grey curve represents the interaction strength among the same segments but when they are disordered. The binding interaction strength difference between disordered and ordered peptides differs by almost an order of magnitude. The same calculations performed using the CHARM36m force field^106^ are shown in Figure S1, where the obtained difference in binding strength between disordered and structured peptides is of the same order.

### Conclusions

In this work, we investigate the impact of enhanced inter-protein interactions in the modulation of the kinetic and thermodynamic properties of ageing biomolecular condensates. Our coarse-grained protein model shows that condensates remain liquid-like when proteins bind to one another weakly (i.e., <1 *k*_B_T), but strengthening of inter-protein interactions (i.e., *>*5 *k*_B_T per residue) gradually slows down the mobility of proteins over time, leading to progressive rigidification/maturation of the condensates^23, 51^. We also observe that aged condensates exhibit a significant degree of hysteresis: once long-lived ordered–ordered interactions are established, amorphous condensates become heat resistant up to moderate temperatures close to the critical conditions for phase separation^110^. Consistently, our atomistic simulations, reveal that formation of inter-peptide *β*-sheets, such as those that may form within the LARKS regions of NUP-98^77, 111^, and similarly in FUS, TDP-43 or hnRNPA1 among other proteins ^23, 68, 77, 112^, can increase the interaction strength between these segments significantly. Such strong binding variation may contribute to rationalise the physicochemical and molecular factors behind the intricate process of pathological maturation and formation of amorphous phase-separated condensates observed in LARKS-containing proteins such as FUS^51^, hnRNPA1^23^, TDP-43^113^, or NUP-98^77, 97^.

We also illustrate how the coupled effects of the decay in protein mobility, the timescale for the emergence of long-lived interactions, droplet coalescence times, and droplet size, crucially govern the shape and material properties of the condensates. When strong inter-protein binding occurs faster than droplet coalescence, the resulting condensates are non-spherical^114, 115^. However, when the strengthening of protein interactions emerge after condensate formation (i.e., once a spherical droplet is already formed), the condensate only experiences a very slight deformation remaining mostly spherical. The time required for two separate tangent droplets to fuse and rearrange into a single spherical condensate depends on both, the initial size of the droplets that are attempting to fuse, and the strength of inter-protein interactions. In small condensates, where the rearrangement time is shorter than the timescale in which proteins lose their mobility due to clustering of structured motifs, condensates are mostly spherical but can eventually become kinetically arrested. In contrast, in larger droplets, where coalescence times are longer, the loss of protein mobility occurs faster than the time required for the condensate to rearrange, and therefore, protein aggregates become kinetically trapped in non-spherical or partially-fused states. Taken together, our results shed light on how local strengthening of inter-protein interactions—for instance due to formation of inter-protein *β*-sheets^77–79, 82, 83^, establishment of post-translational modifications^84^, or changes in salt conditions^26^—may impact the mesoscopic phase behaviour of biomolecular condensates, and suggest a mechanism for the emergence of aspherical droplets over time.

## Acknowledgements

This project has received funding from the European Research Council (ERC) under the European Union Horizon 2020 research and innovation programme (grant agreement No 803326). J.R.E. acknowledges funding from the Oppenheimer Fellowship and from Emmanuel College Roger Ekins Research Fellowship. A.G. acknowledges funding from the EPRSC Doctoral Programme Training number EP/N509620/ and Winton Advanced program. J.A.J. is a Research Fellow at King’s College. This work has been performed using resources provided by the Cambridge Tier-2 system operated by the University of Cambridge Research Computing Service (http://www.hpc.cam.ac.uk) funded by EPSRC Tier-2 capital grant EP/P020259/1.

## Author contributions statement

R.C. G. and J.R.E conceived the project. A.G. and J.R.E conducted the simulations. A.G. and J.R.E analyzed the results. J.A.J. and R.C. G. contextualized the results. A.G. and J.R.E wrote the initial version of the manuscript. All the authors reviewed and edited the manuscript.

