## Supplementary information for "Kinetic interplay between droplet maturation and coalescence modulates shape of aged protein condensates"

(Dated: October 7, 2021)

### SI. ATOMISTIC POTENTIAL OF MEAN FORCE CALCULATIONS

To quantify the interaction strength variation in the protein binding energy due to structural disorder-to-order transitions, we perform atomistic Potential of Mean Force (PMF) simulations [1] of a Low-complexity Aromatic-Rich Kinked Segment (LARKS) of NUP-98 protein [2] before (while remaining disordered) and after undergoing the structural disorder-to-order transition. The employed NUP-98 LARKS sequence was GFGNFGTS (Protein Data Bank (PDB) code 6BZM). The resolved crystalline structure is a $\beta$-sheet domain of four assembled peptides of the same sequence. We perform these simulations in two separate manners: 1) fixing the conformation of the peptide to the one in the PDB crystalline structure (code 6BZM), and 2) allowing them to freely sample their conformational landscape (i.e., as when they are disordered), while keeping the position of the central atom (the one closest to the center of mass of the initial protein conformation, structured protein) constant to control the relative distances between different peptides.

To that end, a set of atomistic (including explicit ions and water) umbrella sampling Molecular Dynamics simulations were carried out using GROMACS 2018 [3] and both a99SB-*disp* [4] and CHARMM36m [5] force fields (results for CHARMM36m simulations will be discussed later). For the a99SB-*disp* [4] force field, we include its model of water and ions which has been proven to work well both with structured and disordered proteins [4]. Since the peptides are relatively short, we add them capping groups: Acetyl to the first backbone nitrogen atom and N-methyl to the last backbone carbon atom.

For the simulations where peptide conformations are kept rigid, the reaction coordinate is the Center of Mass (COM) distance between the dissociating peptide and the one in front of it, whereas in the case of the disordered peptides, since the COM continuously changes due to conformational dynamic rearrangements, we take the distance between the central atoms in each of the chains (these are chosen as the ones closest to the original COM of the chains in the conformation of the crystal). We place a window approximately every 0.3 Å along the reaction coordinate from 0.46 nm until approximately 2.5 nm where longest (electrostatic) range interactions completely vanish (PMFs flat out). To be able to adequately sample a steep potential such as the one found between the rigid structures, we choose a spring constant of 8000 kJ/(mol nm$^2$). Additionally, in the simulations in which we keep the structured peptides rigid, we use positional restraints on the heavy atoms of the chains (excluding hydrogens) in the three directions of space except for the dissociating chain, whose atoms we do not restrain in the direction of the reaction coordinate. We find that restraints of around 1000 kJ/(mol nm$^2$) are enough to keep the molecules rigid. In the case of the disordered peptides, we restrain the positions of the central atoms of each of the chains in the three directions of space except for the dissociating chain whose central atom is not restrained in the direction of the reaction coordinate (pulling atom direction).

For each simulation window, we employ a NaCl concentration of $\sim$ 0.1M. The temperature is set to 300K. The timestep for the integration of the Verlet equations of motion is set to 1 fs. For the rigid chain, we run simulations of about 20 ns (each Umbrella window). Due to the higher number of degrees of freedom of the disordered peptides, each window is run as an independent simulation 5 times to ensure adequate sampling. We run for 25 ns each independent simulation (with a different initial velocity distribution) to complete a total simulated time of 125 ns. To constrain bond lengths and angles we used the LINCS algorithm [6] with an order of 8, and 2 iterations. The cut-off for the Coulombic and Van der Waals interactions has been chosen at a conservative value of 1.4 nm. Particle Mesh Ewald summations for electrostatic interactions are employed [7] with 4th interpolation order and with a fourier spacing of 0.12 nm and Ewald tolerance of 1.5e-5. The simulations are performed in the NpT ensemble, where temperature is kept constant using a Nose-Hoover thermostat [8] with 1 ps relaxation time. To keep pressure constant, we employ a Parrinello-Rahman [9] isotropic barostat at p=1 bar and with a relaxation time of 1 ps.

For each simulation system, we use a box of approximately 12 x 4.2 x 4.2 nm (the long side of the simulation box corresponds to that of the Umbrella Sampling pulling direction). Thus, we typically use around 7234 water molecules and 14 NaCl ion pairs together with 4 protein peptides. All our systems are electroneutral (i.e., the total charge is zero). After solvation of the peptides, we perform an energy minimization with a force tolerance of 1000 kJ mol$^{-1}$ nm$^{-1}$ followed by a short 1000 ps NPT equilibration both with 10000 kJ nm$^{-2}$ mol$^{-1}$ of positional restraints for the heavy atoms of the chains in all three directions of space. The analysis of the simulations was carried out using the WHAM [10] analysis tool implemented in GROMACS [3]. The first 10% time block of the simulations was discarded as equilibration time, although we note that including it barely changes our PMF results.

#### A. PMF calculations with the CHARMM36m force field

Using similar parameters and equivalent setups as those described above for the a99SB-*disp* force field [4], we repeat the simulations with a competing force field, CHARMM36m [5], which can also provide reasonably accurate predictions for folded and disordered proteins [4]. As it can be seen in Fig. S1, while the absolute binding interaction strength before and after the disorder-to-order structural transition is moderately different compared to that predicted by the amber-based force field, the ratio of interactions between structured (blue curve) and disordered (red curve) binding strength is, similarly, of about an order of magnitude (as shown in Fig. 4 of the main text). This result gives us confidence to believe that our calculations for the relative binding interaction strength do not significantly depend on the force field choice, and are a general feature of LARKS structural transitions.

### SII. COARSE-GRAINED IDP MODEL FOR STUDYING STRONG *VS.* WEAK PROTEIN BINDING

#### A. Model description

In our coarse-grained (CG) model for intrisically disordered proteins (IDPs), we describe IDPs as 39-bead polymers of Lennard-Jones beads connected by harmonic springs. The potential energy of our CG force field is given by:

$$E_{Pot} = E_{\text{Bonds}} + E_{\text{Angles}} + E_{\text{LJ}} \tag{S1}$$

where $E_{\text{Bonds}}$ accounts for the bonding energy between two consecutive beads, $E_{\text{Angles}}$ for the associated bond-angle interaction among 3 consecutive beads, and $E_{\text{LJ}}$ corresponds to non-bonded interactions (i.e., interactions between beads that are not directly bonded to each other, either from the same chain or different ones). $E_{\text{Bonds}}$ is described by:

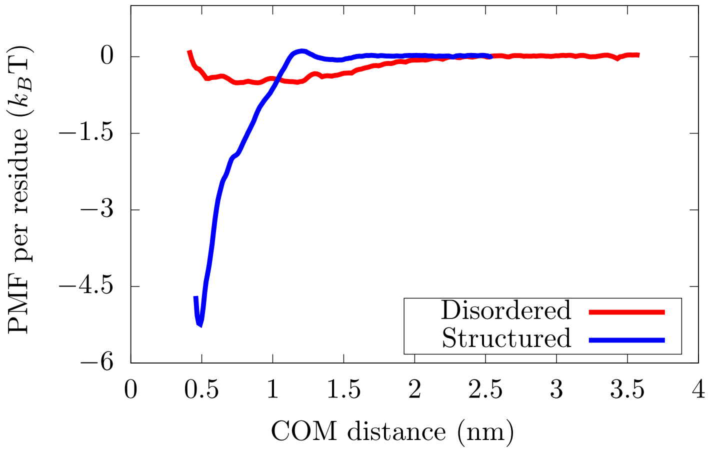


**FIG. S1:** Atomistic Potential of Mean Force (PMF) dissociation curve of an 8-amino acid segment (PDB code: 6BZM) of NUP-98 protein from an aggregate of 4 segments as a function of the center of mass distance (COM) with the CHARM36m force field [5]. Red curve depicts the PMF interaction strength among segments showing a well defined folded structure, kinked $\beta$-sheet structure, while blue curve represents the interaction strength among the same domains but when they are unstructured (fully disordered). Further details of these PMF calculations are provided in Section SI.

$$E_{\text{Bonds}} = \sum_{\text{Protein bonds}} k_{bonds}(r - r_0)^2, \tag{S2}$$

where the distance between distinct bonds is $r$ and the equilibrium bond length is $r_0 = \sigma$. Note that $\sigma$ is the bead molecular diameter and our unit of length in our CG simulations. The spring constant is $k_{bond} = 40\ \epsilon/\sigma^2$, being $\epsilon$ our reduced unit of energy as it will be explained below. The term $E_{\text{Angles}}$ is described by:

$$E_{\text{Angles}} = \sum_{\text{Protein angles}} k_{angle}(\theta - \theta_0)^2, \tag{S3}$$

where $k_{angle} = 3\epsilon/\text{rad}^2$, $\theta$ represents the bond-angle, and $\theta_0$ the equilibrium resting angle ($\theta_0 = 180$). Non-bonded interactions ($E_{\text{LJ}}$) are described by a Lennard-Jones potential:

$$E_{\text{LJ}} = 4\epsilon \left[ \left(\frac{\sigma}{r}\right)^{12} - \left(\frac{\sigma}{r}\right)^{6} \right] \tag{S4}$$

where $r$ accounts for the inter-bead distance, and $\epsilon$ defines the maximum attractive interaction among different beads of a given type. The cut-off for the LJ interactions is set to

2.5 $\sigma$. Our model distinguishes two types of interactions: those belonging to strong 'sticker-sticker' domains where $10\epsilon = \epsilon_S$ (interactions between blue and red beads, as shown in Fig. 1A of the main text), and those belonging to weak 'spacer-spacer' domains, where $\epsilon = \epsilon_D$ (interactions between grey-grey, grey-blue, grey-red, red-red and blue-blue beads). These distinct values of $\epsilon_S$ and $\epsilon_D$ allow the model to effectively consider dramatically different binding interactions between distinct protein domains as those observed in our atomisitic PMF simulations (Fig. S1 and Fig. 4 of the main text).

We have studied two different patterning of our IDP model: one in which strong longer-lived interacting domains are located along the full sequence (Fig. S2A (Top)), and another in which these strongly binding domains are only placed over the first half of the sequence (Fig. S2A (Bottom)). The results for the first sequence are shown in the main text, while those for the second one in Section SII D, although as discussed in the main text and Section SII D, both predict a qualitatively similar ageing behaviour over time.

Moreover, we also test the implications of mapping strong long-lived interactions through the Wang-Frenkel (WF) potential [11] to evaluate the impact of a different coarse-graining approach in the condensate ageing behaviour predicted by our IDP model. In Fig. S2B, we depict in pink the alternative potential curve for coarse-graining enhanced protein binding via the Wang-Frenkel potential [11]. The equation and model parameters of the WF potential leading to the pink curve employed to mimic these interactions in our CG model is given by:

$$u_{WF} = \begin{cases} \alpha\epsilon_S \left[(\frac{\sigma}{r})^2 - 1\right] \left[(\frac{r_c}{r})^2 - 1\right]^2 r \leq r_c \\ 0; \qquad\qquad\qquad\qquad\qquad\qquad \text{if } r > r_c \end{cases} \tag{S5}$$

where $\epsilon_S$ controls the depth of the potential, and it is 10 times larger than $\epsilon_D$, and $r_c$ controls the range of the potential, and it is set to $1.5\sigma$. In the WF potential, $\alpha$ is given by the following expression:

$$\alpha = 2\left(\frac{r_c}{\sigma}\right)^2 \left(\frac{3}{2\left((\frac{r_c}{\sigma})^2 - 1\right)}\right)^3 . \tag{S6}$$

In our coarse-grained simulations, we employ reduced units to refer to the following

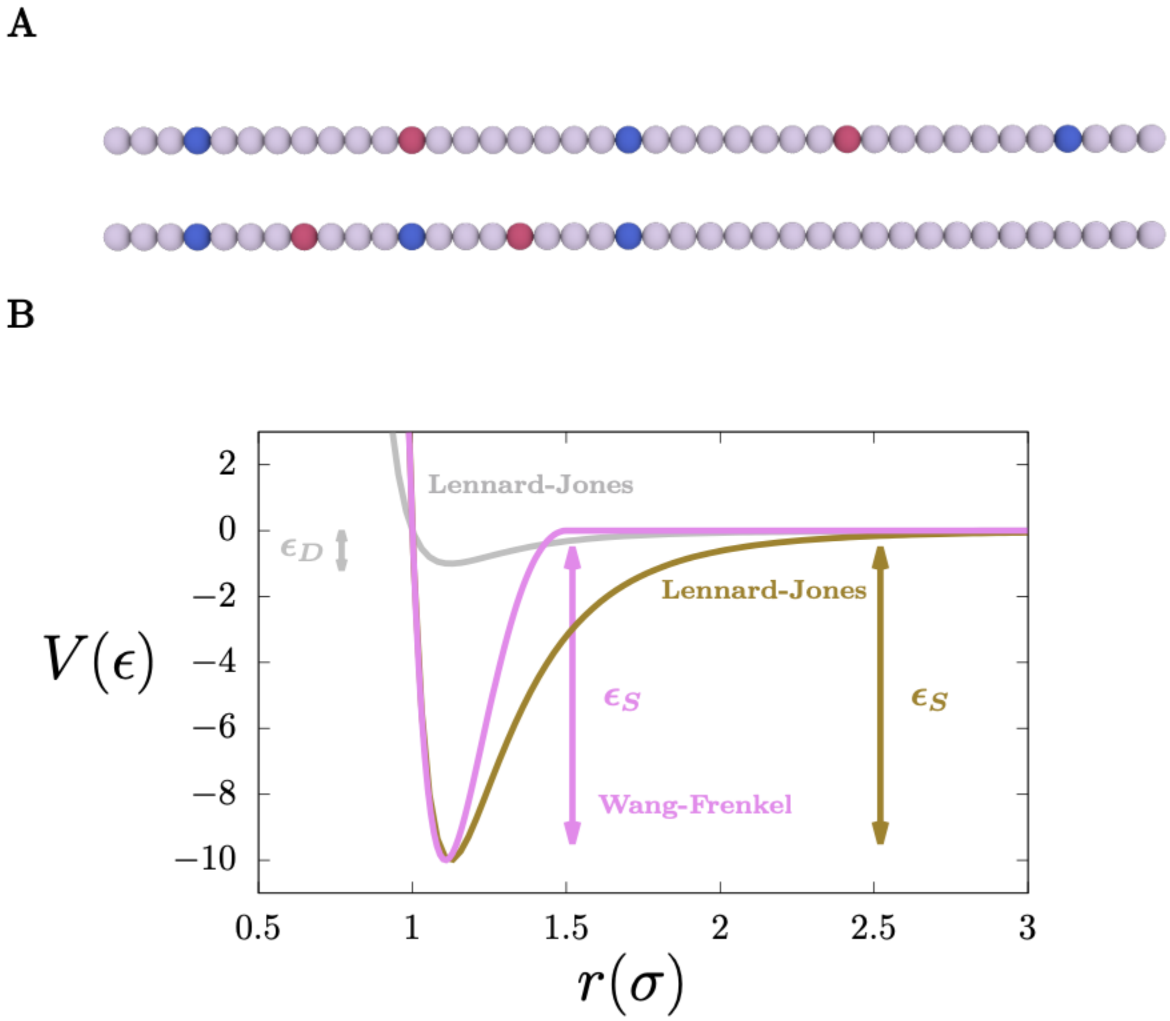


**FIG. S2:** A) Sequence patterning of strong *versus* weak interactions of the two studied IDP coarse-grained models. Red and blue beads account for complementary strong binding motifs, while grey ones for weakly interacting protein regions as described in Fig. 1A of the main text. B) Coarse-grained potential that encodes weak-weak interactions between protein domains (grey curve), and strong long-lived interactions among red and blue protein segments (brown curve). A Lennard-Jones potential of different well-depth ($\epsilon_D$ for interactions between weakly and transient interacting domains, and $\epsilon_S$ for interactions between protein domains with enhanced binding) has been used to model different binding energies between distinct IDP segments in our CG simulations shown in the main text. Moreover, an alternative coarse-graining approach through the Wang-Frenkel potential [11] has been used to model 'sticker-sticker' interactions (pink curve) in order to evaluate the impact of the chosen potential range in mimicking strong binding within our IDP CG approach (see Section SII E).

magnitudes: temperature as $T^* = k_B\mathrm{T}/\epsilon_D$, number density as $\rho^*$ =(N/V)$\sigma^3$, pressure as $p^*$

$= p\sigma^3/\epsilon_D$, and time as $\sigma\sqrt{m/\epsilon_D}$. The mass of all different types of beads is set to $m^* = 1$. We modulate the effective temperature of our simulations by keeping constant T at 300K and varying the interaction strength among IDP beads, $\epsilon/k_B$ (i.e., $\epsilon_S/k_B$ = 1500K=$10\epsilon_D/k_B$, and therefore, $\epsilon_S$=$5k_B$T and $\epsilon_D$=$0.5k_B$T).

#### B. Simulation details

All our coarse-grained simulations have been carried out using LAMMPS [12] software. The integration timestep used to numerically solve the equations of motion was $\Delta t^* = 0.0004$. For NVT and NpT simulations, we employed the Nosé-Hoover thermostat and barostat [8] with relaxation times of $\Delta t^* = 0.4$ and $\Delta t^* = 0.401$ respectively. Except where explicitly mentioned otherwise along the main text (as in coalescence simulations), system sizes were kept constant at 729 IDP replicas, and densities for bulk systems (those used to compute the diffusion coefficient) were equilibrated at the droplet coexistence density of each studied temperature, while for the rest of the simulations (i.e., droplet formation and coalescence simulations) the system total densities were of $\rho^*$ ≈0.01-0.02. In our NVT simulations, to evaluate the time-evolution of the protein diffusion coefficient at bulk droplet conditions, we employ NpT short simulations (of about 10000 time steps) between consecutive NVT production runs to ensure that, despite small changes in the condensate bulk density may occur due to the emergence of strong long-lived contacts, the bulk condensate density corresponds to that of the droplet in equilibrium with the diluted phase at the studied conditions. Please note that these short equilibration NpT simulations have not been considered for the calculation of the mean square displacement to obtain the diffusion coefficient.

#### C. Determination of strong protein contacts

To globally evaluate the number of strong ($s$-$s$) protein interactions in a given system as a function of time, we compute the specific interaction energy between red and blue beads, and we divide it by the depth of the $s$-$s$ interaction potential ($\epsilon_S$). On the other hand, to locally identify the regions of the condensate that exhibit $s$-$s$ contacts, we also use a local order parameter which individually determines the number of IDP red and blue beads that are engaged to other strongly binding domains for a given cut-off distance. Protein domains with enhanced binding (red or blue beads) which are in contact with at least 3 other complementary bead pairs in a cut-off distance of $2.1\sigma$ are considered to be part of a region of high-density long-lived interactions with low protein mobility (as depicted in green in the spherical condensates time-evolution of Fig. 2B of the main text).

#### D. Comparison of different IDP sequence patterning in condensate maturation

In this section, we report the ageing behaviour of condensates formed by IDPs that possess the sequence patterning of strongly *vs.* weakly interacting domains depicted in Fig. S2A (Bottom) while keeping the same LJ potential employed in the main text (brown curve of Fig. S2B) to mimic (*s*-*s*) interactions. In this patterning, segments with strong binding are only distributed over the first half of the sequence (Fig. S2A (Bottom)).

In Fig. S3, we show the dependence of the diffusion coefficient ($D$) as a function of time for IDPs within phase-separated condensates at different protein interactions strengths (similarly to Fig. 1B of the main text but for the sequence patterning shown in Fig. S2A (Bottom)). As it can be seen, for 'sticker-sticker' interactions with binding energy $\epsilon_S \geq 6 k_B$T, protein diffusion slows down over time indicating gradual condensate maturation. On the contrary, for lower binding *s*-*s* interactions (i.e., $\epsilon_S$=5$k_B$T), liquid-like behaviour persists over time with no hints of protein mobility deceleration (orange curve). Nevertheless, we note that $D$ for this patterning at $\epsilon_S$=5$k_B$T is moderately lower than that at the same $\epsilon_S$ for the sequence patterning shown in Fig. S2A (Top), and Fig. 1B of the main text. Snapshots of IDP spontaneous condensation along time in the liquid-like regime (Fig. S3 Top Right) and ageing regime (Fig. S3 Bottom Right) are also included. Even though the enhanced binding energy threshold separating liquid-like behaviour from gradual kinetic arrest moderately depends on the IDP patterning ($\Delta\epsilon_S$ can vary $\sim 1 k_B$T), the same qualitative behaviour shown in Fig. 1B of the main text is recovered here.

Moreover, we investigate whether aged condensates formed by this IDP sequence also exhibit thermal hysteresis upon maturation. To that purpose, as in Fig. 2A of the main

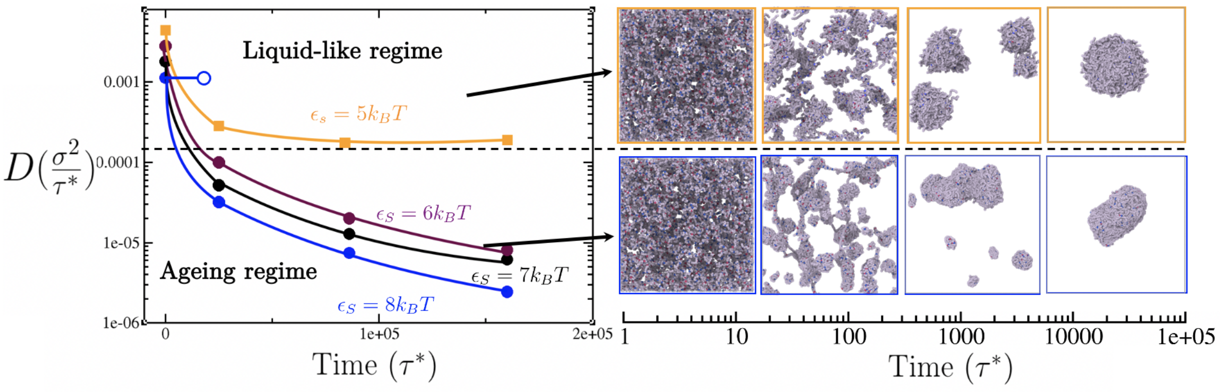


**FIG. S3:** Time-evolution of the IDP diffusion coefficient ($D$) in the condensed phase for different interaction strengths $\epsilon_S$ (in $k_B$T) between strongly-binding protein segments. The IDP sequence patterning corresponds to that shown in Fig. S2A (Bottom). The same coarse-graining approach (LJ potential) to model 'sticker-sticker' enhanced protein binding employed in the main text is also used here. The horizontal black dashed line represents the kinetic threshold of our simulation timescale that distinguishes between ergodic liquid-like behaviour and ageing (transient liquid-to-solid) behaviour. Interaction strengths lower than $6k_{\rm B}$T between segments displaying strong binding permit liquid-like behaviour (up to $\epsilon_S = 3.5k_{\rm B}$T and $\epsilon_D = 0.35k_{\rm B}$T where LLPS is no longer possible), while equal or higher $s$-$s$ binding lead to the gradual deceleration of protein mobility over time as shown by $D$. However, in absence of strongly-binding segments, where all beads bind to one another with uniform binding strength, liquid-like behaviour can be still observed even at $\epsilon_D$ values of $0.8k_{\rm B}$T (empty blue circle) denoting the key role of enhanced protein binding in gradual condensate rigidification. Black arrows indicate the time dependent behaviour of condensates over time in the liquid-like (Top) and ageing regimes (Bottom). The time evolution snapshots of the condensate corresponds to systems with $\epsilon_S = 5k_{\rm B}$T (Top) and $\epsilon_S = 7k_{\rm B}$T (Bottom). Please note that these snapshots do not correspond to the NVT bulk systems employed to compute the diffusion coefficient in the Left panel.

text, we perform two simulations starting from two different initial configurations: 1) from a homogeneous single phase (Fig. S4 Top), and 2) from a matured amorphous condensate formed under ageing conditions (Fig. S4 Bottom). As it can be seen, even though the conditions of both simulations are the same (i.e., protein-protein interactions are $\epsilon_S$=5$k_B$T and $\epsilon_D$=0.5$k_B$T), the shape of the condensates over time is different (spherical and non-spherical respectively), thus, suggesting thermal hysteresis as also observed for the patterning illustrated in Fig. S2A (Top) and shown in Fig. 2B of the main text. Therefore, we acknowledge that patterning effects on IDP sequences exhibiting long-lived *vs.* weak/transient contacts, even leading to a similar qualitative ageing behaviour, can have a significant impact on the required thermodynamic conditions to observe thermal hysteresis as well as in the threshold of 'sticker-sticker' interaction strength to switch condensates from the liquid-like behaviour to the maturation regime as shown in Figs. S3 and S4 (i.e., protein diffusion coefficient).

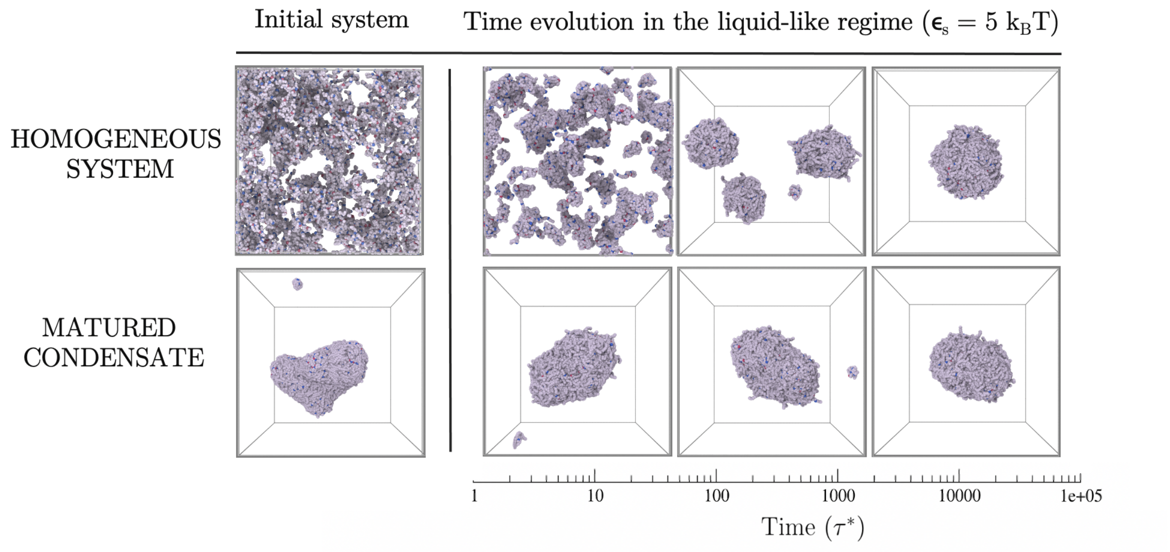


**FIG. S4:** Thermal hysteresis of the condensates probed via coarse-grained simulations with the IDP sequence patterning shown in Fig. S2A (Bottom) and the same IDP coarse-graining LJ potential used in the main text (brown and grey curves of Fig. S2B). Top panel: Time-evolution starting from an homogeneous system where protein–protein interactions are moderate (i.e., $\epsilon_S$= $5k_B$T and $\epsilon_D$= $0.5k_B$T). Bottom panel: Time-evolution at the same conditions above, although starting from a matured condensate that was formed under strong ageing conditions ('sticker-sticker' protein interactions of $\epsilon_S$= $8k_B$T).

#### E. Comparison of distinct coarse-graining approaches in modelling enhanced protein binding interactions

In this section, we report the ageing behaviour of condensates formed by IDPs that possess the sequence patterning of strongly *vs.* weakly interacting domains depicted in Fig. S2A (Top) (the same as in the main text) but using the WF potential to model enhanced binding interactions (pink curve in Fig. S2B).

In Fig. S5, we plot the dependence of the diffusion coefficient as a function of time for IDPs within phase-separated condensates at different protein interactions strengths (as in Fig. 1B of the main text but using the WF potential described in Eq. S5). As it can be seen, for 'sticker-sticker' binding interactions of $\epsilon_S \geq 6k_B$T, protein diffusion slows down over time indicating gradual condensate rigidification. On the other hand, for lower interactions (i.e., $\epsilon_S$=5$k_B$T), liquid-like behaviour persists over time with no hints of protein mobility deceleration (orange curve). Snapshots of IDP spontaneous condensation along time in the liquid-like regime (Fig. S5 Top Right) and ageing regime (Fig. S5 Bottom Right) are also included. Even though the *s*-*s* binding energy threshold separating the liquid-like behaviour from the gradual kinetic arrest moderately depends on the coarse-graining potential and interaction range of the enhanced binding motifs ($\Delta\epsilon_S \sim +0.7k_B$T compared to the same patterning with LJ interactions for strongly-binding motifs), the same qualitative behaviour shown in Fig. 1B of the main text is recovered here.

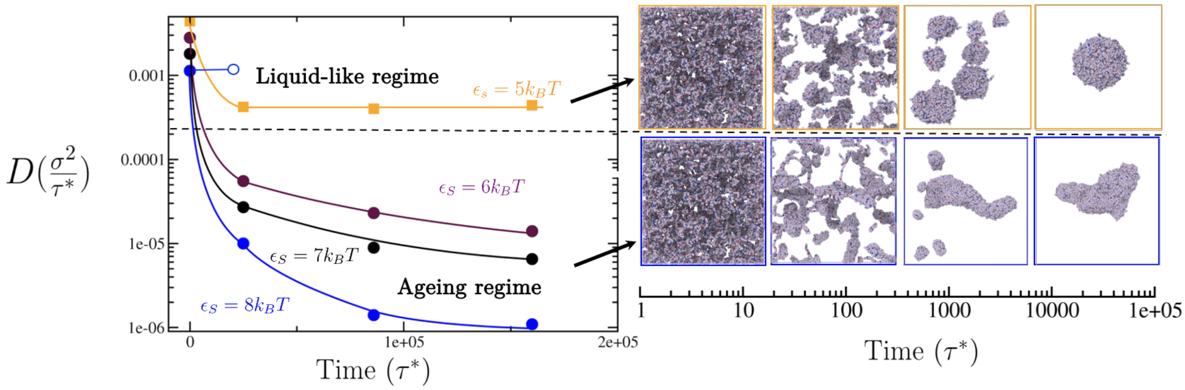


**FIG. S5:** Time-evolution of the IDP diffusion coefficient ($D$) in the condensed phase for different interaction strengths $\epsilon_S$ (in $k_B$T) between strongly-binding protein segments. The IDP sequence patterning corresponds to that shown in Fig. S2A (Top) (as in the main text), but using the Wang-Frenkel potential [11] to model strong binding as shown in Fig. S2B (pink curve). The horizontal black dashed line represents the kinetic threshold of our simulation timescale that distinguishes between ergodic liquid-like behaviour and ageing (transient liquid-to-solid) behaviour. Interaction strengths lower than 6$k_\mathrm{B}$T between strongly-binding segments permit liquid-like behaviour (up to $\epsilon_S = 3.5k_\mathrm{B}$T and $\epsilon_D = 0.35k_\mathrm{B}$T where LLPS is no longer possible), while equal or higher *s*-*s* binding strengths lead to the gradual deceleration of protein mobility over time as shown by $D$. However, in absence of strongly-binding interactions, where all beads bind to one another with uniform binding strength, liquid-like behaviour can be still observed even at $\epsilon_D$ values of 0.8$k_\mathrm{B}$T (empty blue circle) denoting the key role of strong binding motifs in gradual condensate rigidification. Black arrows indicate the time dependent behaviour of condensates over time in the liquid-like (Right Top) and ageing regimes (Right Bottom). The time evolution snapshots of the condensate corresponds to systems with $\epsilon_S = 5k_\mathrm{B}$T (Top) and $\epsilon_S = 7k_\mathrm{B}$T (Bottom). Please note that these snapshots do not correspond to the NVT bulk systems employed to compute the diffusion coefficient time-evolution shown in the Left panel.

Moreover, we investigate whether aged condensates in which strong binding (e.g., due to

LARKS ordered-ordered interactions) is modelled through the Wang-Frenkel potential also exhibit thermal hysteresis upon maturation. To that purpose, as in Fig. 2A of the main text and Fig. S4, we perform two simulations starting from two different initial configurations: 1) from a homogeneous single phase (Fig. S6 Top), and 2) from a matured amorphous condensate formed under ageing conditions (Fig. S6 Bottom). As it can be seen, even though the conditions of both simulations are the same (i.e., protein-protein interactions correspond to $\epsilon_S$=5.8$k_B$T and $\epsilon_D$=0.58$k_B$T), the shape of the condensates over time is completely different (spherical and non-spherical respectively), thus, suggesting thermal hysteresis as observed for the same IDP patterning, but using the LJ potential as shown in Fig. 2A of the main text. Therefore, even though we acknowledge that the coarse-graining approach to describe enhanced peptide interactions can have a moderate impact on the required thermodynamic conditions to observe thermal hysteresis ($\Delta\epsilon_S \sim 0.8k_B$T) as well as in the threshold of 'sticker-sticker' interaction strength to switch from liquid-like behaviour to transient rigidification (maturation regime), the same qualitative maturation behaviour is observed independently of the chosen potential (Figs. S5 and S6).

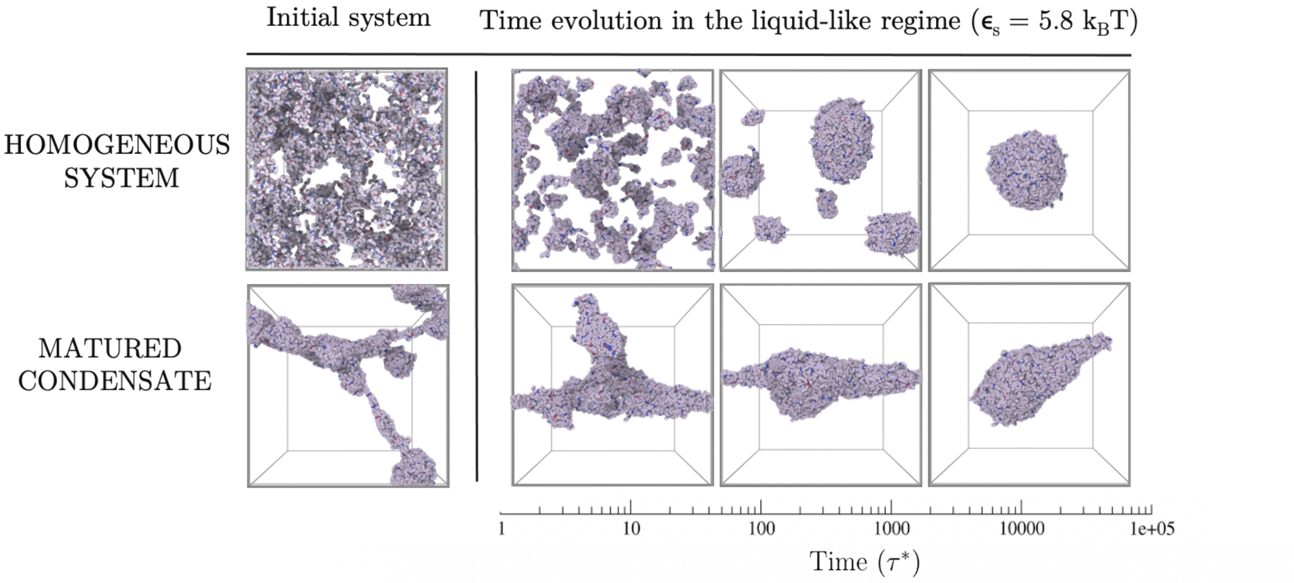


**FIG. S6:** Thermal hysteresis of the condensates probed via coarse-grained simulations with the IDP sequence patterning shown in Fig. S2A (Top) but using the Wang-Frenkel potential [11] to model strongly-binding interactions as shown in Fig. S2B (pink curve). Top panel: Time-evolution starting from an homogeneous system where protein–protein interactions are moderate (i.e., $\epsilon_S$= 5.8$k_B$T and $\epsilon_D$= 0.58$k_B$T). Bottom panel: Time-evolution at the same conditions above, although starting from a matured condensate that was formed under strong ageing conditions ('sticker-sticker' protein interactions of $\epsilon_S$= 8$k_B$T). Please note that even though partial rearrangement of the aged condensate can occur at $\epsilon_S$= 5.8$k_B$T for longer windows of time than those in which spherical condensates emerge from the homogeneous single phase (Top), matured condensates remain mostly kinetically trapped (in non-spherical shapes) over the studied windows of time.
